# Catch and kill airborne SARS-CoV-2 to control spread of COVID-19 by a heated air disinfection system

**DOI:** 10.1101/2020.06.13.150243

**Authors:** Luo Yu, Garrett K. Peel, Faisal H. Cheema, William S. Lawrence, Natalya Bukreyeva, Christopher W. Jinks, Jennifer E. Peel, Johnny W. Peterson, Slobodan Paessler, Monzer Hourani, Zhifeng Ren

## Abstract

Airborne transmission of severe acute respiratory syndrome coronavirus 2 (SARS-CoV-2) *via* air-conditioning systems poses a significant threat for the continued escalation of the current coronavirus disease (COVID-19) pandemic. Considering that SARS-CoV-2 cannot tolerate temperatures above 70 °C, here we designed and fabricated efficient air disinfection systems based on heated nickel (Ni) foam to catch and kill SARS-CoV-2. Virus test results revealed that 99.8% of the aerosolized SARS-CoV-2 was caught and killed by a single pass through a Ni-foam-based filter when heated up to 200 °C. Additionally, the same filter was also used to catch and kill 99.9% of *Bacillus anthracis*, an airborne spore. This study paves the way for preventing transmission of SARS-CoV-2 and other highly infectious airborne agents in closed environments.

**One Sentence Summary:** Heated Ni-foam filters are capable of effectively catching and killing airborne SARS-CoV-2 and *Bacillus anthracis* spores.

## Main Text

Coronavirus disease (COVID-19), caused by the severe acute respiratory syndrome coronavirus 2 (SARS-CoV-2), is a rapidly spreading pandemic that is severely threatening public health all over the world (*1–6*). According to the World Health Organization (WHO), as of June 11, 2020, there have been more than 7.4 million confirmed cases in 216 countries, areas, or territories, leading to at least 418,294 deaths (*7*). The rapid spread of COVID-19 is related to SARS-CoV-2 carriers being highly infectious while asymptomatic and the high capability of the virus to survive in various environmental conditions (*8–10*). The most probable SARS-CoV-2 transmission route is human-to-human (*11–13*), which may explain why cluster spread was the main reason for the quickly increasing cases of COVID-19 in early February 2020 in Wuhan, China (*14*). Additionally, the consensus among scientists is that the virus is also noticeable when it is transmitted through aerosols and droplets that are released into the air by a carrier, especially when the person coughs, sneezes, or even talks forcefully in a closed environment (*15, 16*). A recent study compared simulated SARS-CoV-2 aerosols to those of SARS-CoV-1 (*17*), its most closely related viral strain and the cause of the 2003 SARS outbreak in Asia, and showed that, comparable to the case of SARS-CoV-1, aerosols containing SARS-CoV-2 can remain in the air for about 3 hours, although their viral load continually diminishes during that time. In the same study it was also found that the virus contained in droplets that settled on various surfaces can remain viable for several days.

Having confirmed airborne transmission of SARS-CoV-2, a mode of transmission that is relevant to several previously known respiratory viruses, including influenza, scientists are now questioning whether the virus can travel even greater distances through the air by becoming lodged in other airborne particles such as condensed water vapor or even dust. Such a mode of transmission would be extremely concerning and would call into question the adequacy of measures that are mostly designed to address issues related to proximity to an infectious individual, such as wearing masks, washing hands and surfaces, and general social distancing. One of the earliest studies addressing this subject indicated that such transmission may be possible for SARS-CoV-2 (*18*). The study looked at certain indicators of airborne viral spread in a Wuhan hospital where patients with COVID-19 were kept in isolation. With all available precautions in place to prevent viral spread through personnel or equipment, viral RNA was still detected in areas of the hospital that it could only have reached through the atmosphere or the ventilation system (*18*). Currently, with increasing numbers of people returning to the workplace, the chances of infection resulting from aerosol transmission through central air-conditioning systems are increasing. Thus, determining how to stop the virus from spreading in air-conditioned spaces is extremely urgent. Simple filtration cannot completely stop the spread. Fortunately, most viruses, including SARS-CoV-2, are not resistant to high temperature (*19, 20*). It has been demonstrated that the time needed for SARS-CoV-2 inactivation is reduced to 5 minutes when the incubation temperature is increased to 70 °C (*21*). Therefore, if a filter in an air conditioner can be heated to a high temperature (*e.g*., up to 250 °C), any SARS-CoV-2 in the cycling air can be efficiently killed in a very short time. An even more challenging task is to prevent the transmission of other airborne highly infectious agents that have been used for bioterrorism, such as *Bacillus anthracis* (anthrax) spores. *Bacillus anthracis* is a large (1-1.5 μm × 3-10 μm in size), aerobic, gram-positive, spore that has long been considered a biological warfare agent (*22*).

Traditional air-conditioner filters based on fiberglass or aluminum (Al) mesh are difficult to heat or have large pores (about 1 centimeter in size), so they cannot effectively catch and kill the virus contained in aerosols (generally smaller than 5 μm in size) (*23*) or other airborne highly infectious agents, such as anthrax spores. An ideal filter should be self-heated rather than have an external heat source that would surely cause a very large rise in air temperature, which requires that the filter itself be electrically conductive. Commercial nickel (Ni) foam is electrically conductive and mechanically strong and it exhibits good flexibility, properties that have prompted its wide use in energy conversion and storage applications (*24–26*). More importantly, Ni foam is highly porous with randomly located pores that are between 50 and 500 μm in size and that meander from one side of the foam to the other, resulting in a very large surface area that can effectively catch particles in the air passing through the filter due to van der Waals forces. The self-heated filter has the additional advantage that the heating is localized on the Ni foam and heat transfer to the passing air is minimal due to the short time of contact between the air and the Ni foam and also the very low thermal conductivity of air (0.02 W m^-1^ K^-1^). Therefore, Ni foam may act as a good filter for catching and killing SARS-CoV-2 or anthrax spores in air-conditioning systems. However, it is extremely challenging to design such a filter since the resistivity of Ni foam is too small to achieve heating at a sufficiently high temperature. In order to realize a filter for preventing the spread of SARS-CoV-2 and anthrax spores, here we designed and fabricated a filter device consisting of folded pieces of Ni foam in multiple compartments connected electrically in series to efficiently increase the resistance to a manageable level so that a temperature up to 250 *°C* was able to be achieved, and found that the filter device exhibits almost 100% ability to catch and kill aerosolized SARS-CoV-2 and anthrax spores in air passed once through the Ni foam heated up to 200 *°C* (temperature optimization will be addressed in a future study). Our study demonstrates the possibility of applying commercial Ni foam as an air-conditioner filter for use in airplanes, airports, hospitals, schools, office buildings, restaurants, hotels, cruise ships, *etc*. for 100% removal of SARS-CoV-2 in cycling air, thus slowing the spread of COVID-19, as well as to prevent transmission of other airborne highly infectious agents like anthrax spores.

The optical image in Fig. 1A shows that commercial Ni foam has typical metal luster, and it is highly flexible, so it can be easily molded into different shapes like the accordion folds shown in Fig. 1B. Due to its high porosity of ≥95% (Fig. S1A), Ni foam also exhibits very high air permeability, as indicated by the clear observation of light passing through Ni foam under the glare of a fluorescent lamp (Fig. 1C). The Ni foam pore size is in the range of ~50 to 500 μm, and the diameter of a single Ni wire is about 65 μm, as seen in the scanning electron microscope (SEM) images in Figs. 1D, E, respectively. The cross-section SEM (Fig. 1F) and optical (Fig. S1B) images further reveal that the thickness of the Ni foam is around 1.6 mm and that the Ni wires in the foam are randomly interconnected with one another, creating a three-dimensional (3D) network structure with many non-straight channels. Therefore, although the aerosols containing SARS-CoV-2 or anthrax spores are smaller than the pores in the Ni foam, it remains highly probable that the aerosolized SARS-CoV-2 and anthrax spores will be captured by the heated Ni wires due to the meandering path, in contrast to the straight path resulting from the well-organized wires in an Al mesh. It should be noted that the thickness, pore size, and porosity of Ni foam can all be easily controlled during the manufacturing process in case different pore sizes are required for different sizes of viruses or other infectious agents. The X-ray diffraction (XRD) pattern in Fig. 1G demonstrates the pure phase of Ni with strong diffraction peaks from the (111), (200), and (220) planes. We then calculated the electrical resistivity (ρ) of a strip of commercial Ni foam 250 mm × 10 mm × 1.6 mm in size. From the slope of the current (I)-voltage (V) curve shown in Fig. 1H, the resistance (R) of the Ni foam is about 0.178 Ω at near room temperature, so the ρ is calculated to be around 1.42 × 10^-5^ Ω m, indicating good conductivity. Thus, we can electrically heat a piece of Ni foam to the required temperature under a certain input power. As shown in Fig. 1I, the temperature of the Ni foam increases very quickly under increasing input power, and it reaches to about 119 *°C* when the input power is ~6.6 W.

**Fig. 1.**
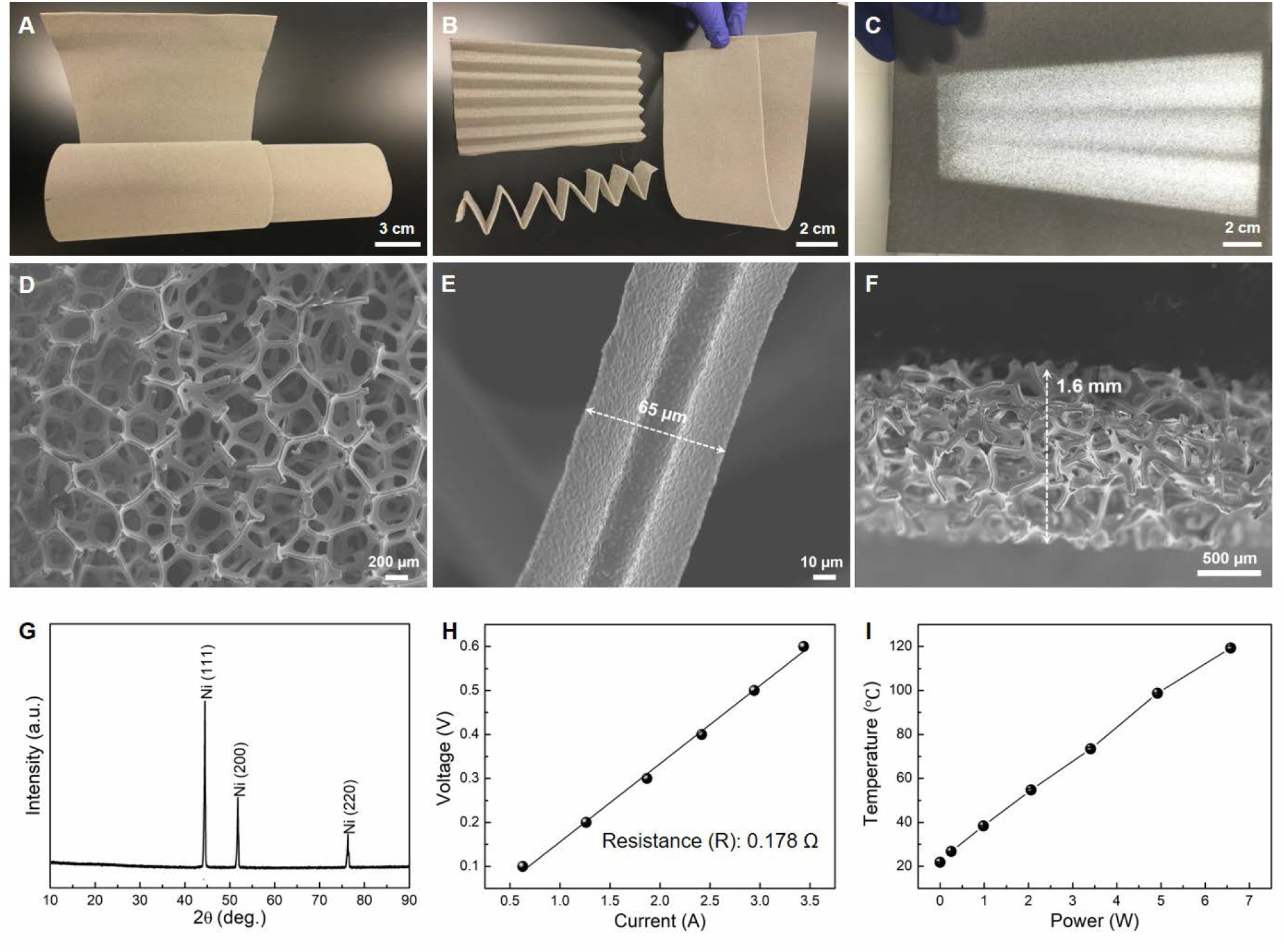
Basic properties of commercial Ni foam. (**A-C**) Photographs under different conditions. Photograph (**C**) was taken under the glare of a fluorescent lamp. (**D, E**) Surface SEM images at different magnifications. (**F**) Cross-section SEM image. (**G**) XRD pattern. (**H**) I-V curve of a strip of Ni foam 1.6 mm × 250 mm × 10 mm in size. (**I**) T-P curve showing the relationship between the Ni foam temperature and the input power.

For use in air conditioners, we must also consider the temperature of the air after it passes through the heated Ni foam; if the air temperature becomes too high, the heated Ni foam would not be suitable for large-scale application. To test this, we blew high-purity nitrogen (N2) gas toward one side of the heated Ni foam and measured the air temperature at different distances away from its opposite side. The distance between the N_2_ gas source and the heated Ni foam was ~3.5 cm and the room temperature was about 21.7 °C. From the results shown in Fig. S2, we can see that the air temperature decreases very quickly after passing through the heated Ni foam. Even for the Ni foam with a high temperature of 115.3 °C, the air temperature is close to room temperature at 4 cm away. Therefore, we do not need to be concerned about the air temperature becoming too high after passing through the hot Ni-foam filter. It should be noted that the resulting air temperature inside ductwork may be slightly higher than in an open environment as we tested.

Due to the very low resistivity of Ni foam, it is not possible to simply use a single piece of Ni foam as a filter that satisfies both the size requirement for heating, ventilation, and air conditioning (HVAC) systems and the U.S. residential voltage requirement (110 V). Considering the flexibility of Ni foam, we designed a folded structure (Fig. 2A) that exhibits much larger resistance due to its significantly increased length, ideally addressing both of the above requirements. In addition, compared with a flat Ni-foam filter, the folded one has two other advantages. First, as illustrated in Fig. 2B, if the thickness of the Ni foam is 1.6 mm, the distance for catching and killing viruses or other infectious agents is only 1.6 mm when it is flat. However, after folding, the effective distance can be much longer, *e.g*., 10 times that for the flat Ni foam if considering a bending length of 1.6 cm, since the gaps within the folds retain sufficiently high temperature to kill viruses and other infectious agents efficiently. It should be noted that the bending length can be easily controlled, and the longer the bending length, the higher the temperature. Second, compared to the flat Ni foam with two main sides exposed to the air, the folded Ni foam has a much smaller surface area exposed to the incoming and outgoing air, which minimizes the heat loss so that the temperature of the Ni foam increases much more quickly and can reach a much higher value at the same power consumption. As shown in Fig. 2C, under the same voltage of 1.0 V, the temperature of the folded Ni foam is 114.9 °C, which is more than twice that of the flat Ni foam (55.8 °C).

Finally, we fabricated filters (Fig. 2D) that each use six pieces of folded Ni foam connected electrically in series, which effectively increases the total resistance to a manageable level so that regular-gauge electrical wires can be used. To enhance the efficiency for catching and killing SARS-CoV-2 and anthrax spores, two filters were parallelly assembled inside a closed device (Fig. S3). We first studied the I-V and temperature (T)-resistance (R) curves for the filter. As the results in Fig. 2E show, the Ni foam filter exhibits a typical metal property, in which the resistance increases with increasing temperature. We further investigated the influence of air flow on the temperature of the filter, and the results are shown in Fig. 2F. Clearly, with an air flow rate of 10 L min^-1^, the temperature of the filter shows a slight decrease of about 10 *°C* under the same input power relative to that without air flow.

**Fig. 2.**
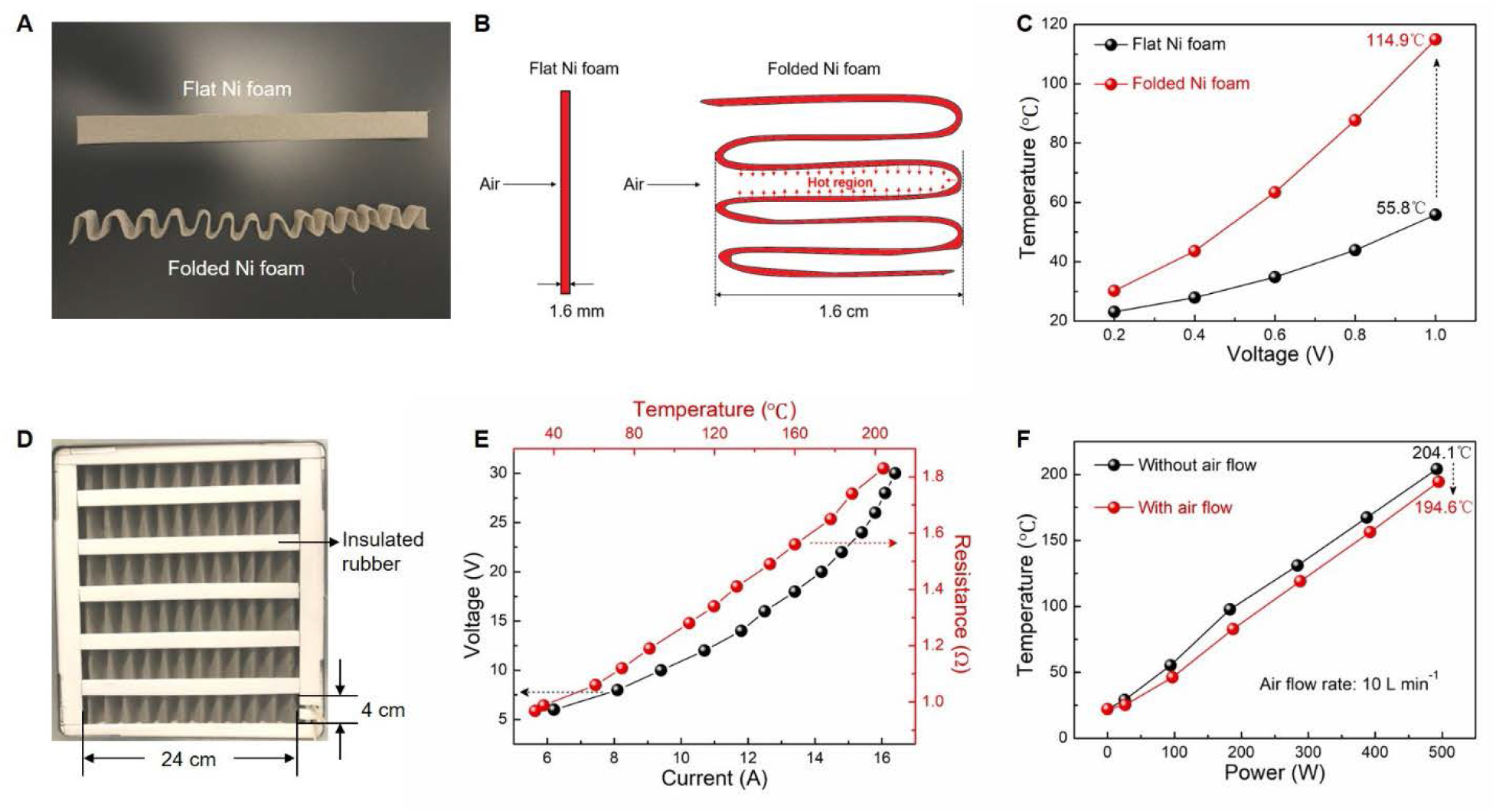
Study of Ni foam as a filter. (**A**) Photographs and (**B**) side-view schematic illustrations of flat Ni foam and folded Ni foam. (**C**) Temperature comparison between flat Ni foam and folded Ni foam under the same voltage. The Ni foam is 20 mm × 250 mm × 1.6 mm in size, and the bending length is about 1.6 cm. (**D**) Photograph of the fabricated filter using six pieces of folded Ni foam connected electrically in series. (**E**) I-V and T-R curves of the filter. (**F**) T-P curves of the filter with and without air flow (high-purity N_2_ gas).

The prototype device was tested using aerosolized actual SARS-CoV-2, isolated from humans, and encouragingly demonstrated 99.8% viral load reduction from upstream to downstream in the device using a single passthrough when the filter was heated up to 200 °C (temperature optimization is currently being studied). As shown in Fig. 3A, by using the median tissue culture infectious dose (TCID50) method for determining viral titer reduction, we confirmed that, compared to the control without a filter, the heated filter significantly reduced the viral titers of SARS-CoV-2 through a single pass in this prototype device. A 2.7-fold log reduction was noted when the Ni-foam filter was heated to ~200 °C (Fig. 3B). Note that the reduction shown by the unheated filters is also effective but the virus is not killed by the unheated filters, which poses a health threat to those who later change the filters. Therefore, the preferred mode is to heat the filter so that the virus is completely caught and killed, leaving no potential future concern. After the successful demonstration of catching and killing almost 100% of the SARS-CoV-2, we designed similar experiments for testing the elimination of *Bacillus anthracis* (anthrax spores) in a separate prototype device. The heated Ni-foam filter was found to catch and kill 99.9% of the anthrax spores through a single pass in the prototype device (Fig. 3A) and a 3.23-fold log reduction (Fig. 3C) was achieved when the Ni-foam filter was heated.

**Fig. 3.**
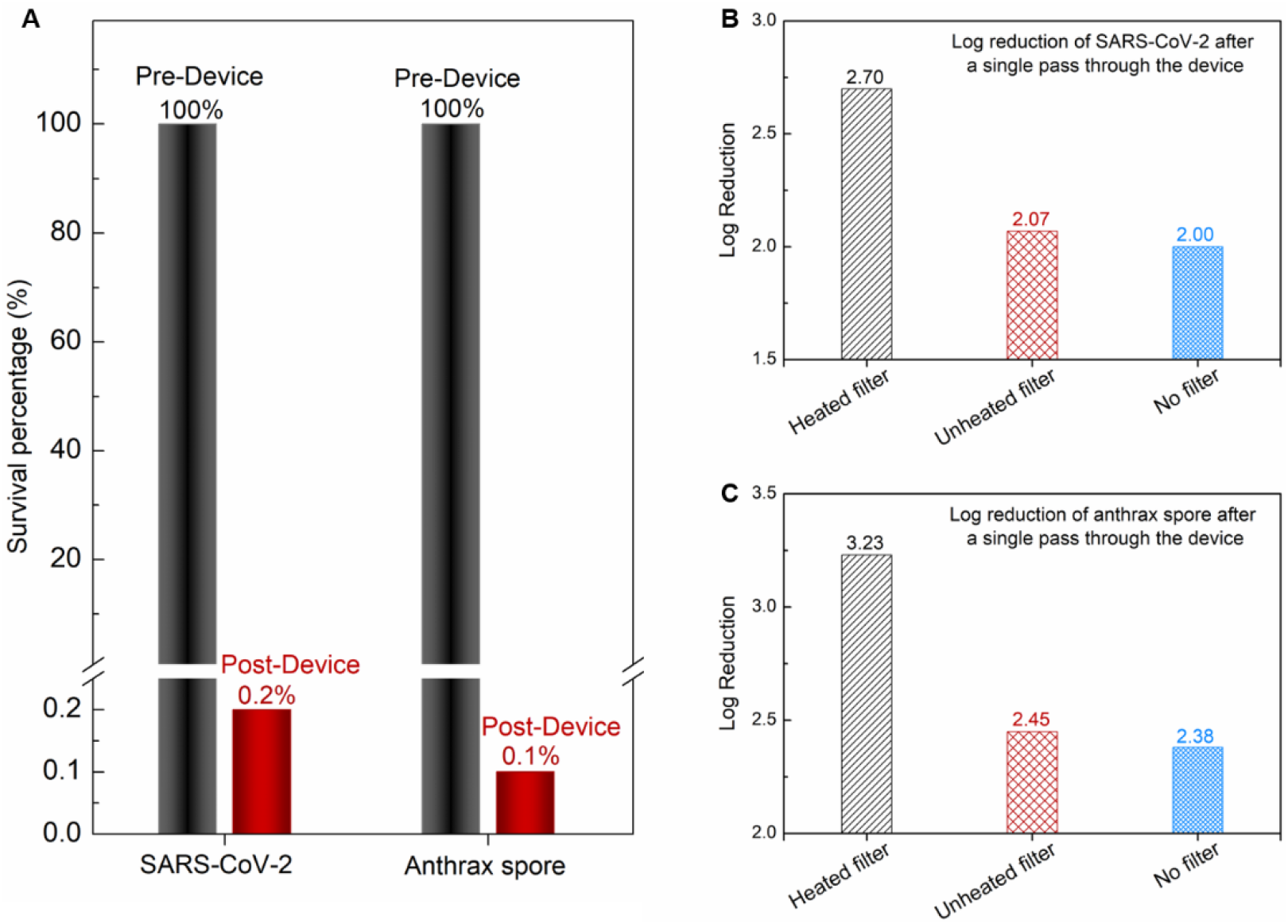
Performance of prototype device on aerosolized SARS-CoV-2 and *Bacillus anthracis*. **(A)** Absolute reduction of TCID50 of aerosolized SARS-CoV-2 and *Bacillus anthracis* by heated filters, showing 99.8% and 99.9% reductions, respectively, between pre-device and post-device levels. Log reduction by the heated filter, unheated filter, and control (No filter) for (**B**) SARS-CoV-2 and (**C**) *Bacillus anthracis*.

The deployment of these novel filter and purification units stands to have a dramatic impact on both essential workers and the general public in the current COVID-19 pandemic, as well as reducing the risk of exposure to other airborne highly infectious agents, both known and unknown. With a phased roll-out, beginning with high-priority venues where essential workers are at elevated risk of exposure (particularly hospitals and healthcare facilities, as well as public transit environs such as airplanes), this innovative technology will (a) improve the safety for frontline workers in essential industries by minimizing the risk of SARS-CoV-2 exposure, (b) make it possible for non-essential workers to safely return to public work spaces by reducing their risk of exposure, and (c) allow for the general public to more safely re-engage with their own communities through the creation of mobile air-purification devices that can be carried on one’s person in order to maintain clean personal air space. These outcomes will enable resilience in the battle against COVID-19, in which the front lines are everywhere and rapidly changing. This technology will also provide for safe bioagent protection gear to eliminate future bioterrorism threats from airborne infectious agents such as anthrax. The air purification and disinfection system derived from this Ni-foam-based heated filter, in conjunction with ultraviolet germicidal irradiation, will be a useful addition in the armamentarium of technologies available to combat future pandemics.

## Author contributions

M.H. and G.K.P. conceived the filter concept for catching and killing the airborne viruses. Z.R. and L.Y. conceived the electrically conducting porous filter idea. C.J. made the testing systems. Z.R., L.Y., G.K.P., J.W.P, S.P. and F.H.C. designed the experiments. L.Y. performed all the temperature and power measuring experiments and collected the data. W.S.L., J.E.P., and N.B. conducted the virus and spore experiments and analyzed the data. L.Y., Z.R., G.K.P. and F.H.C. analyzed all the data and results, and wrote the paper. All authors contributed to the discussion of the results and commenting on the manuscript.

## Data and materials availability

All data are available in the manuscript and supplementary materials.

## Supplementary Materials

Materials and Methods

Figures S1-S3

