## Supplementary Materials for "Catch and kill airborne SARS-CoV-2 to control spread of COVID-19 by a heated air disinfection system"

**This PDF file includes:**

Materials and Methods

Figs. S1-S3

### Materials and Methods

Ni foam

Pieces of commercial Ni foam were purchased from MTI Corporation in USA and Kunshan Jiayisheng Electronics Co Ltd in China. Both have very similar properties in terms of purity (>99.99%), surface density (346 g $m^{-2}$), thickness (1.6 mm), and porosity (≥95%). The pieces of Ni foam were used without any treatment.

Morphology and nanostructure

The surface morphology and nanostructure of the Ni foam were examined by scanning electron microscopy (SEM, JEOL JSM-6330F).

Phase composition

Phase composition of the Ni foam was characterized by X-ray diffraction (PANalytical X'pert PRO diffractometer).

Current and temperature measurements

We used a programmable DC power supply (BK PRECISION, 9153, 60 V/9A) to measure the voltage (V)-current (I) curves and to heat the Ni-foam filters. The temperature was recorded using an infrared thermometer (MICRO-EPSILON, CTL-CF2-C3).

Aerosolization method

Aerosolization of SARS-CoV-2 and *Bacillus anthracis* Ames spores was accomplished using an automated aerosol control platform (Biaera AeroMP; Biaera Technologies, LLC). The bioaerosols were generated using either a 3-jet (*B. anthracis* spores) or a 6-jet (SARS-CoV-2) Collison nebulizer, both of which typically produce droplet sizes of approximately 1 μm. The total flow rate to the filtration unit, comprised of nebulizer air and diluter air, was 30 L $min^{-1}$. The nebulizer air flow rates to the 3- and 6-jet Collison nebulizers were 7.5 and 14 L $min^{-1}$, respectively. Bioaerosol samples were collected before and after filtration for each aerosol run using S.K.C.

BioSamplers (S.K.C., Inc.). The flow rate to each BioSampler was approximately 10 L $min^{-1}$. Aerosolization and bioaerosol sampling was performed for either 15 min (*B. anthracis* spores) or 20 min (SARS-CoV-2). All aerosolization procedures were performed in a Class III biosafety cabinet housed within the animal biosafety level 3 (ABSL-3) facility of the Galveston National Laboratory (GNL).

<u>Virus titration determination method</u>

Vero cells (ATCC Cat# CCL-81, RRID:CVCL_0059) were cultured in Dulbecco's Modified Eagle Medium (DMEM) (HyClone) supplemented with 10% fetal bovine serum and 1% penicillin/streptomycin (HyClone). SARS-CoV-2 USA-WA1/2020 was obtained from the World Reference Center for Emerging Viruses and Arboviruses (WRCEVA), which obtained the original isolate from the Centers for Disease Control and Prevention (CDC). Upon obtaining the isolate, virus titration was performed. A subsequent low multiplicity of infection (MOI) passage was performed in Vero cells (DMEM supplemented with 5% fetal bovine serum and 1% penicillin/streptomycin), and the resulting stock was titrated *via* TCID50 (median tissue culture infectious dose) in Vero cells to determine the titer prior to use.

The 96-well plate with Vero cells to be ~85-95% confluent in 24 hours was prepared. Cells were plated at 2 x $10^5$ $mL^{-1}$. Growth media used was DMEM supplemented with 10% fetal bovine serum and 1% penicillin/streptomycin. The virus sample was diluted in quadruplicate at a 1:10 ratio in DMEM + 5% fetal bovine serum and 1% penicillin/streptomycin, for a total of 6 dilutions ($10^{-1}$ to $10^{-6}$). Media was removed from the cells and replaced with 100 uL of the diluted virus. Cells were incubated at 37 °C and 5% $CO_2$ for 4 days. On day 4, the post-infection virus inoculum was removed from the plate and the cells were fixed with 100 uL of 10% buffered formalin per well. After 30 min of fixation, the formalin was removed and the cells were stained with 0.25% crystal violet. SARS-CoV-2 positive cells were noted and TCID50 was calculated. These experiments were conducted within approved biosafety level three (BSL-3) laboratories at the University of Texas Medical Branch (UTMB) and the GNL.

Anthrax spore production and quantitation

*B. anthracis* Ames spores were grown in modified Schaeffer's medium using a computer-controlled New Brunswick B-510 20-L fermentor operating within a Baker BioProtect II biosafety cabinet installed in the GNL BSL-3 Enhanced Facility. After inoculation, the fermentor was operated with aeration at pH 7.0-7.5 with pH control for approximately 4 days, after which time the crude spore content of the culture was aseptically harvested by centrifugation at 9,000 x g and washed with sterile molecular grade water. The spores were purified by density gradient centrifugation using sterile MD-76. Visual observation of the spores at 400x by phase-contrast microscopy during each step of purification was performed to ensure production of a homogeneous suspension of highly refractile spores.

The bacterial concentration of the samples was determined using an automatic serial diluter and plater (easySpiral Dilute; Interscience). The samples, diluted in sterile water, were plated onto trypticase soy agar plates containing 5% sterile sheep blood (TSAB) and incubated at 37 °C for 16-24 hours. Colonies from the plates were then enumerated using an automatic colony counter (Scan 500; Interscience).

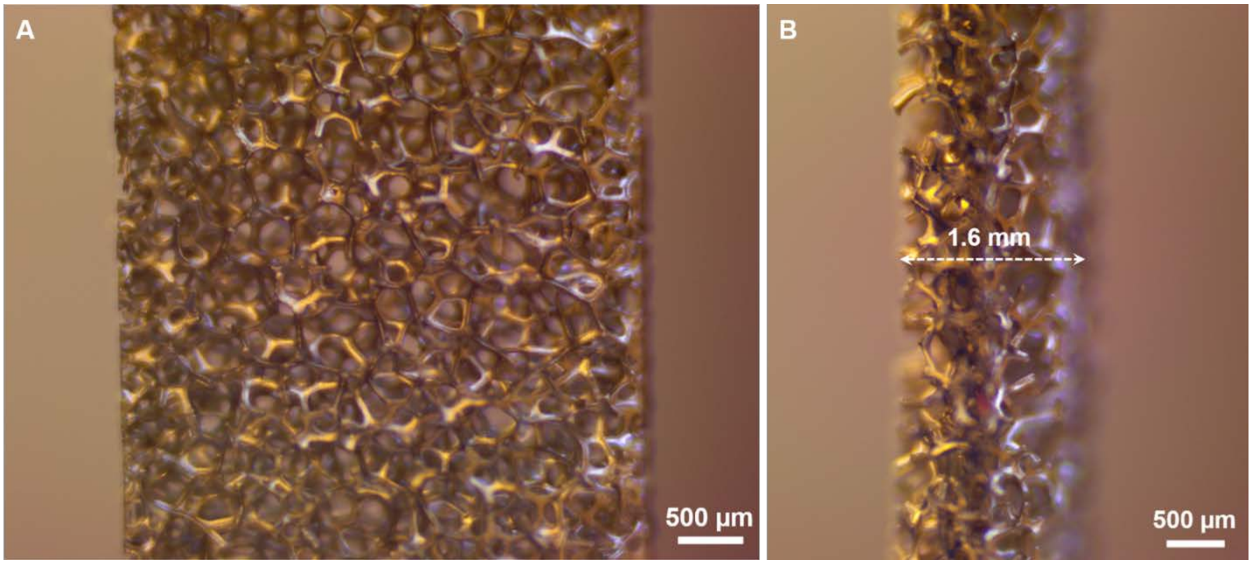


**Fig. S1. Optical images of commercial Ni foam. (A)** Front view and **(B)** side view.

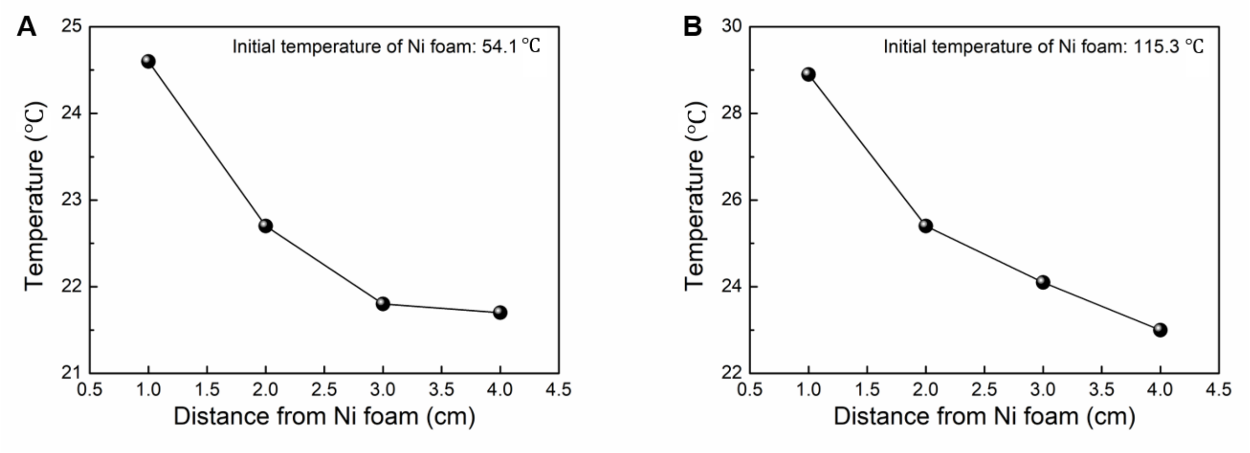


**Fig. S2. Decreasing temperature of air after flowing through heated Ni foam.** Initial Ni-foam temperature of **(A)** 54.1 ℃ and **(B)** 115.3 ℃.

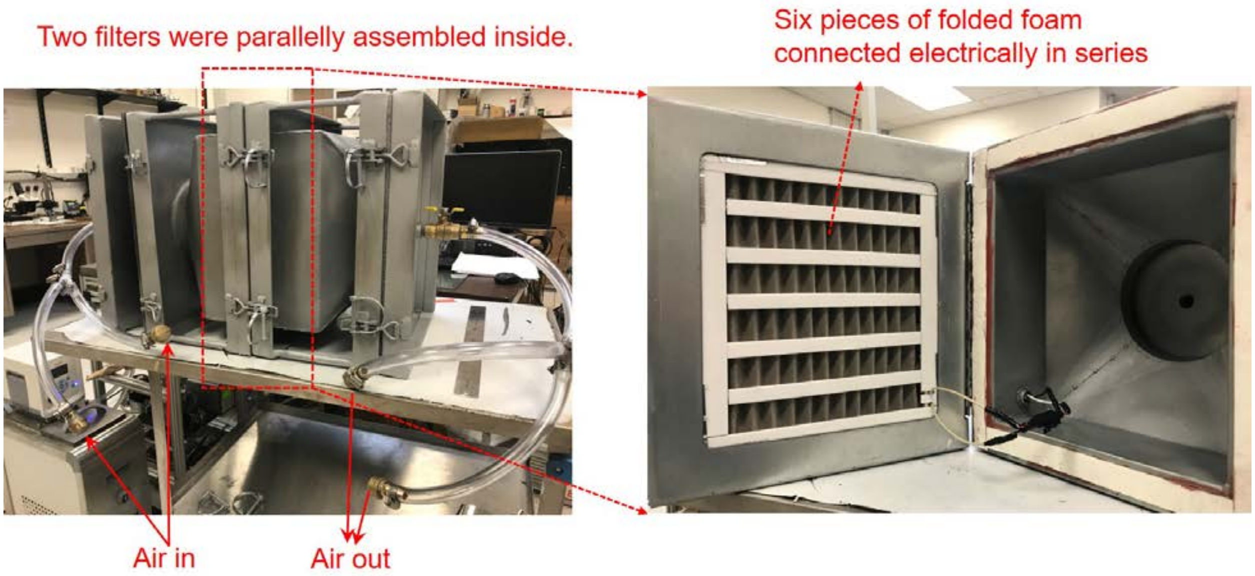


**Fig. S3. Device for virus experiment.** Photographs show the device details (left) and one of the two filters using folded Ni foam (right).
